# Modeling liquid-liquid phase separation and its impact on proteasomal substrate degradation

**DOI:** 10.1101/2025.07.20.665740

**Authors:** Di Wu, Qi Ouyang, Hongli Wang

## Abstract

The proteasome, central to ubiquitin-proteasome mediated degradation, undergoes liquid-liquid phase separation (LLPS) during substrate processing. However, the mechanisms governing proteasomal LLPS and its functional interplay with degradation dynamics remain elusive. Here, we developed a reaction-diffusion model with thermodynamic constraints to quantitatively characterize proteasome-associated LLPS. The model successfully captures experimental LLPS dynamics, revealing a concentration-dependent phase transition boundary influenced by proteasome activity and molecular interactions. Significantly, LLPS was found to have bidirectional (enhancing or suppressing), context-dependent effects on degradation efficiency. This work elucidates regulatory mechanisms and functional roles of LLPS in proteasome-mediated degradation, offering a predictive quantitative model and testable hypotheses for future investigation.

## 1. Introduction

The proteasome, a ∼2.5 MDa macromolecular protease complex, serves as the central degradation machinery of the ubiquitin-proteasome system (UPS) in eukaryotic cells and plays a pivotal role in cellular proteostasis [1,2]. Structurally, it comprises two primary subcomplexes: the regulatory particle (RP), responsible for substrate recognition, deubiquitination, and unfolding, and the core particle (CP), which houses the proteolytic active sites [3,4]. Ubiquitinated substrates bind to the RP via ubiquitin receptors, undergo ATP-dependent unfolding by RP-associated ATPases, and are translocated into the CP for degradation [3,5–7]. Given the intricate dynamics and multi-layered regulation governing proteasome function, extensive experimental and theoretical efforts have investigated its mechanisms across scales, encompassing substrate translocation energetics [3,5–7], substrate selectivity [8–10], RP-CP communication [11–13], and modulation by proteasome-associated proteins [14–17].

Recent studies have revealed a novel layer of proteasome regulation: the formation of proteasomal liquid-liquid phase separation (LLPS) condensates during substrate degradation [18–21]. This phenomenon indicates that beyond the intrinsic dynamics of individual proteasomes, collective interactions between proteasomes, substrates, and auxiliary factors represent a critical aspect of proteasome biology in specific contexts. Initial observations identified RAD23B as a key driver of proteasomal LLPS condensates induced by hyperosmotic stress, with these droplets dissolving as substrates were degraded [19]. Subsequent work confirmed RAD23B-mediated proteasomal LLPS under amino acid starvation [20]. Furthermore, the scaffolding protein p62 has also been shown to nucleate proteasomal LLPS condensates through mechanisms conceptually similar to RAD23B [21–23].

Despite these significant experimental advances, fundamental questions regarding the mechanisms and functional consequences of proteasomal LLPS remain unresolved. While key molecular players (e.g., RAD23B, p62) have been identified [19–22], a quantitative understanding of the physicochemical principles governing condensate formation, stability, and dissolution during active degradation is still lacking. The concept of “UPS overload” has been proposed, suggesting that LLPS arises as a response to excessive proteolytic demand [24]. However, whether a well-defined boundary exists between functional (non-saturated) and overloaded (saturated) states of the UPS remains unclear. Critically, the functional impact of proteasomal LLPS on substrate degradation efficiency – whether it enhances, impedes, or contextually modulates degradation – is a pivotal question for elucidating its biological significance. While theoretical frameworks, primarily based on mean-field theory, have provided valuable insights into general LLPS droplet dynamics (e.g., chemically driven division and stabilization) [25,26], dedicated computational models specifically addressing proteasomal LLPS are needed to bridge this gap and complement experimental approaches.

To address these critical questions, we developed a minimal mechanistic model based on partial differential equations, comprising eight key components derived from current experimental evidence. This model quantitatively characterizes the spatiotemporal dynamics of LLPS during proteasomal substrate degradation. Our simulations recapitulate the experimentally observed formation and subsequent dissolution of condensates over time. Crucially, increasing initial substrate concentration revealed a sharp transition boundary for LLPS occurrence, providing a quantitative definition for the hypothesized UPS saturation state. Model analysis demonstrated that weakening molecular interactions (e.g., substrate valency) or reducing proteasomal degradation rates can shift this saturation boundary, either preventing or promoting the system’s entry into the saturated state. Strikingly, comparing degradation kinetics with and without LLPS revealed that condensate formation can exert bidirectional effects, either promoting or inhibiting overall substrate degradation efficiency depending on system parameters. This study provides novel mechanistic insights into the regulation and functional duality of proteasomal LLPS and generates experimentally testable predictions to guide future exploration of this emerging phenomenon in proteostasis.

## 2 Thermodynamic model of proteasomal LLPS during substrate degradation

The model of RAD23B-mediated LLPS during proteasomal degradation is primarily based on the recent experimental findings [19]. The interaction between substrate and RAD23B and proteasomal reactions for the LLPS-mediated substrate degradation are depicted in Fig. 1. Under hyperosmotic stress, the ubiquitinated substrates (polypeptides) undergo LLPS via interactions between ubiquitin chains and the ubiquitin-associated (UBA) domains of RAD23B. Within the liquid droplets, substrates recruit proteasomes by binding to ubiquitin receptors on the proteasome’s regulatory particle. Proteasomes unfold and degrade substrates within the core particle, with ubiquitin chains progressively removed. Following complete degradation of a substrate, the proteasome binds to new substrates within the droplet, enabling cyclic degradation. Concurrently, substrate degradation reduces substrate concentration. This weakens RAD23B-ubiquitin chain interactions and ultimately leads to droplet dissolution.

**Fig. 1.**
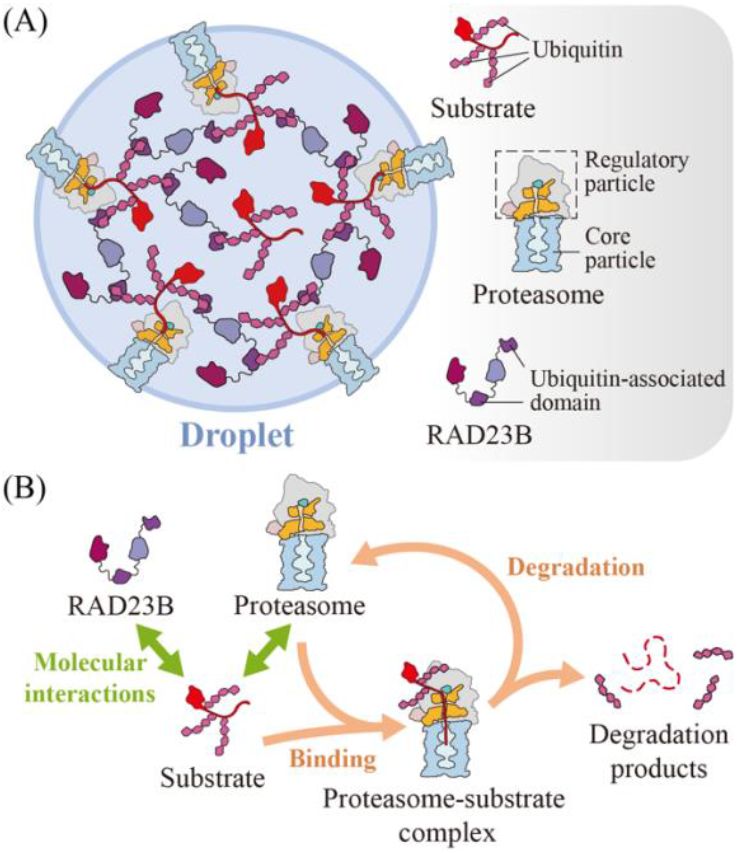
Interactions and reactions involved in proteasomal LLPS. (A) Schematic diagram of proteasomal LLPS and its components. (B) Intermolecular interactions (green arrows) between the components and substrate degradation reactions (orange arrows) in the proteasome.

In our model, we neglect the distinct physicochemical properties of individual substrates and their degradation products, as well as the heterogeneity introduced by other proteins within the cellular environment. The process of substrate degradation in proteasomes is delicate which involves complex proteasomal conformation transitions. Here we do not delve into the detailed changes of proteasomal structures, but adopt a coarse-grained description. For simplicity, we consider only three substrate-proteasome complexes: *PS*_0_, *PS*_1_, and *PS*_2_. Here, *PS*_0_ represents the initial substrate-proteasome binding complex. *PS*_1_ and *PS*_2_ represent subsequent intermediate degradation states characterized by the progressive removal of ubiquitin tags by the proteasome, with *PS*_2_ having undergone more extensive deubiquitination than *PS*_1_. The proteasomal substrate degradation can be represented by the following four reactions,

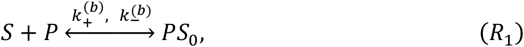

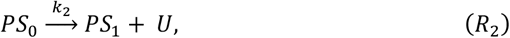

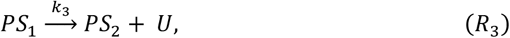

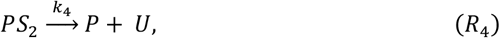

where *S, P*, and *U* denote the substrate, the proteasome, and ubiquitinated peptide fragments (degradation products), respectively. We do not use the simple mass action law to describe the chemical kinetics since phase separation requires non-ideal fluids. Instead, the reaction rates *k*_2_-*k*_4_ in *R*_2_-*R*_4_ are assumed to follow the Arrhenius law, 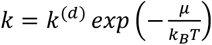, where *μ* is the chemical potential for *PS*_0_, *PS*_1_ and *PS*_2_. Taking into account RAD23B (*R*) and the solvent (*W*), the model includes a total of eight components: *C* = {*R, S, P, U, W, PS*_0_, *PS*_1_, *PS*_2_}.

The RAD23B-mediated proteasomal LLPS is transient during substrate degradation, with the liquid droplets forming at high substrate concentrations and dissolving at low substrate concentrations [19]. This phase separation forms droplets that not only compartmentalize substrate degradations but are also dynamically altered by the proteasomal degradation processes. The intertwined physics of phase separation and chemical reactions in non-ideal solution has been recently reviewed in [27, 28]. To model the proteasomal LLPS and its impact on substrate degradation, we adopt a reaction-diffusion framework described by partial differential equations of the following form,

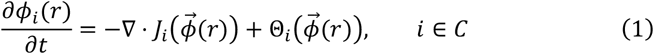

where *ϕ*_*i*_ represents the spatial dependent volume fraction of component *i*, and *C* = {*R, S, P, U, W, PS*_0_, *PS*_1_, *PS*_2_}. 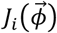 is the flux of component *i* as a function of the volume fractions of all components 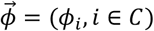, and Θ_*i*_ denotes the changing rate of *ϕ*_*i*_ due to the reactions *R*_1_-*R*_4_. The flux *J*_*i*_ is driven by the gradients of chemical potentials *μ*_*j*_, *j* ∈ *C*, which takes the form,

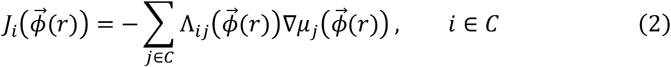

where 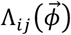 denotes the weight that the gradient Δ*μ*_*j*_ contributes to flux *J*_*i*_. Under the constraint of total zero flux Σ_*i*∈*C*_ *J*_*i*_ = 0 and the normalization condition Σ_*i*∈*C*_ *ϕ*_*i*_ = 1, we assume 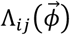 as,

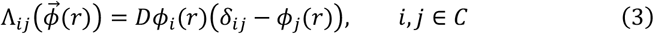

where *D* is the coefficient. The chemical potential *μ*_*i*_ can be writen as [27],

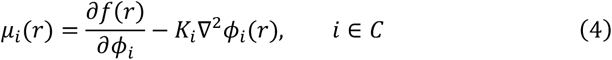

where *f* is the free energy, whose explicit form depends on all the components involved in the proteasomal LLPS and substrate degradation that we considered here. *K*_*i*_ denotes the interfacial tension coefficient for component *i*.

The spatially extended free energy *f* is written as [27],

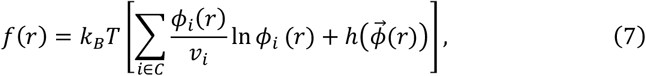

where *v*_*i*_ is the specific volume of component *i* and 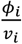 In *ϕ*_*i*_ represents the mixing entropy of component *i*. The enthalpy density function 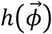 can be expressed as:

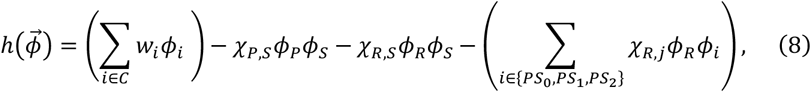

where *w*_*i*_ and *Χ*_*ij*_ are coefficients. In the righthand side of Eq. (8), the second term represents *P-S* interactions, the third term describes *S-R* interactions, and the final term characterizes interactions among *R* and the proteasome-substrate complexes (*PS*_0_, *PS*_1_, *PS*_2_). The substrate *S* maintains identical ubiquitination levels as *S*-bound proteasome *PS*_0_, while *PS*_0_, *PS*_1_, *PS*_2_ have progressively decreasing ubiquitin tags. We assume that 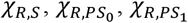 and 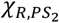 satisfy 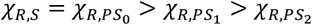 as ubiquitin is removed during substrate degradation, the interaction between the substrate and RAD23B weakens.

The reaction rates Θ_*i*_ ‘s in Eq. (1) can be obtained readily from the binding and degradation reactions *R*_1_*-R*_4_. As there are no chemical reactions for RAD23B and solvent molecules, Θ_*R*_ and Θ_*W*_ are zeros. For the rest of the components *S, P, U, PS*_0_, *PS*_1_, *PS*_2_, the rates Θ_*i*_’s take the following form,

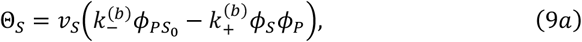

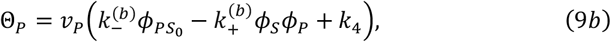

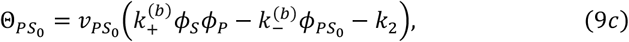

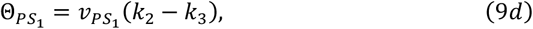

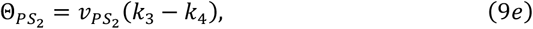

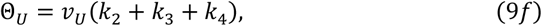

The combination of Eqs. (1-9) represents a physicochemical description for the tight interplay between LLPS and proteasomal substrate degradations with thermodynamic constraints.

## 3. Results

### 3.1 Temporal evolution of proteasomal LLPS

The partial differential equation system Eqs. (1), equipped with Eqs. (2-9), was solved numerically using a finite difference method in both one-dimensional (1D) and two-dimensional (2D) domains. 1D domain size of 48 with a uniform discreteness of 128 points and 2D domain size of 48×48 with a uniform 128×128 grid was simulated with Neumann boundary conditions. Unless otherwise specified, the parameters used in the numerical simulations throughout this work were set to: 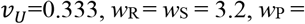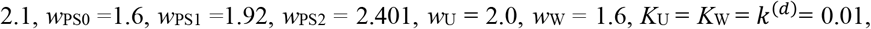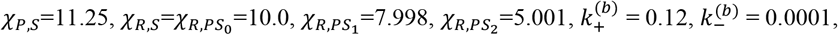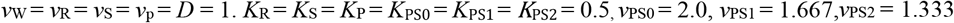 and with homogeneous initial conditions: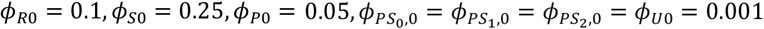. As shown in Fig. 2, the model successfully reproduced LLPS condensation in both 1D and 2D spatial domains, where droplets emerges from initially homogeneous distributions of component. Figure 2A depicts the spatiotemporal evolution in 1D space. Droplets nucleate rapidly, with concentrated component molecules, and subsequently undergo gradual dissolution over time. Within these droplets, distinct spatial segregation is observed as depicted in Figs. 2A and 2C: *PS*_0_ predominantly localizes to the droplet core, while *PS*_1_ and *PS*_2_ accumulate at the droplet edge, with *PS*_2_ distributed closer to the periphery than *PS*_1_. In 2D simulations (Fig. 2B), the model behavior closely parallels the 1D results. *R*/*S*/*P*-enriched droplets form initially and then diminish in number as the proteasome-mediated substrate degradation processes, ultimately nearing complete dissolution. This is consistent with experimental observations [19]. The characteristic spatial distributions of *PS*_0_, *PS*_*1*_ and *PS*_2_ observed in 1D are also recapitulated in the 2D simulations. Figure 2D depicts the typical spatial distributions in the droplet, *i*.*e*., center-distributed (such as *R, S, P, PS*_0_) and edge-distributed (such as *PS*_1_, *PS*_2_) components. The distribution feature originates from the sequential reduction in ubiquitin tags on the proteasome-substrate complexes (*PS*_0_, *PS*_1_, *PS*_2_), which progressively weakens their interaction with RAD23B (*R*). Consequently, complexes exhibiting stronger interactions with *R* (higher ubiquitination) localize inward, while those with weaker interactions (lower ubiquitination) localize outward.

**Fig. 2.**
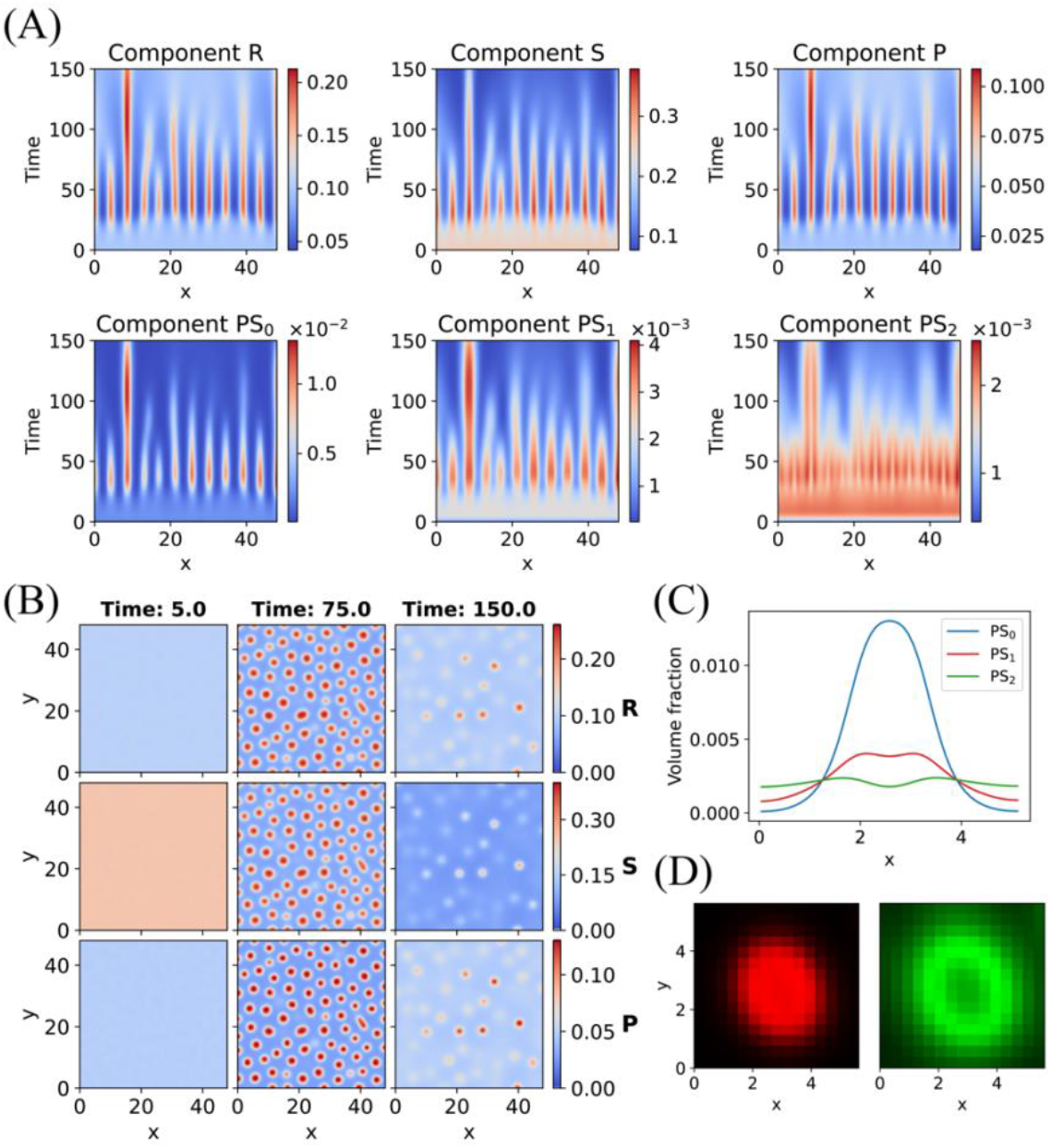
Time evolution of proteasomal LLPS during substrate degradation. (A) Spatiotemporal evolution of LLPS in 1D space for RAD23B (*R*), substrate (*S*), and proteasomes with and without bounded substrates *P, PS*_0_, *PS*_1_ and *PS*_2_. (B) Snapshots of the temporal evolution of LLPS in 2D space for components *R, S*, and *P* at time *t* = 5, 75, 150. (C) Spatial distributions of *PS*_0_, *PS*_1_ and *PS*_2_ in a 1D droplet taken from (A). (D) Two typical spatial distributions in the droplet: center-distributed (red, for *R, S, P* and *PS*_0_) and edge-distributed (green, for *PS*_1_ and *PS*_2_) components.

To further compare with experimental observations, we analyzed the temporal evolution of droplet count and median droplet diameter in our 2D simulations. As shown in Fig. 3, the droplet count increases rapidly to an initial peak, followed by a gradual decrease as substrates are continuously degraded by proteasomes, ultimately leading to the dissolution of droplets. The median droplet diameter also exhibits rapid initial growth, attaining its maximum value near-simultaneously with the peak droplet count. However, in contrast to the droplet count, the median diameter decreases more gradually after reaching its maximum. These simulation results show qualitative agreement with experimental observations [19], indicating that the model captures the essential features of RAD23B-mediated LLPS dynamics during substrate degradation.

**Fig. 3.**
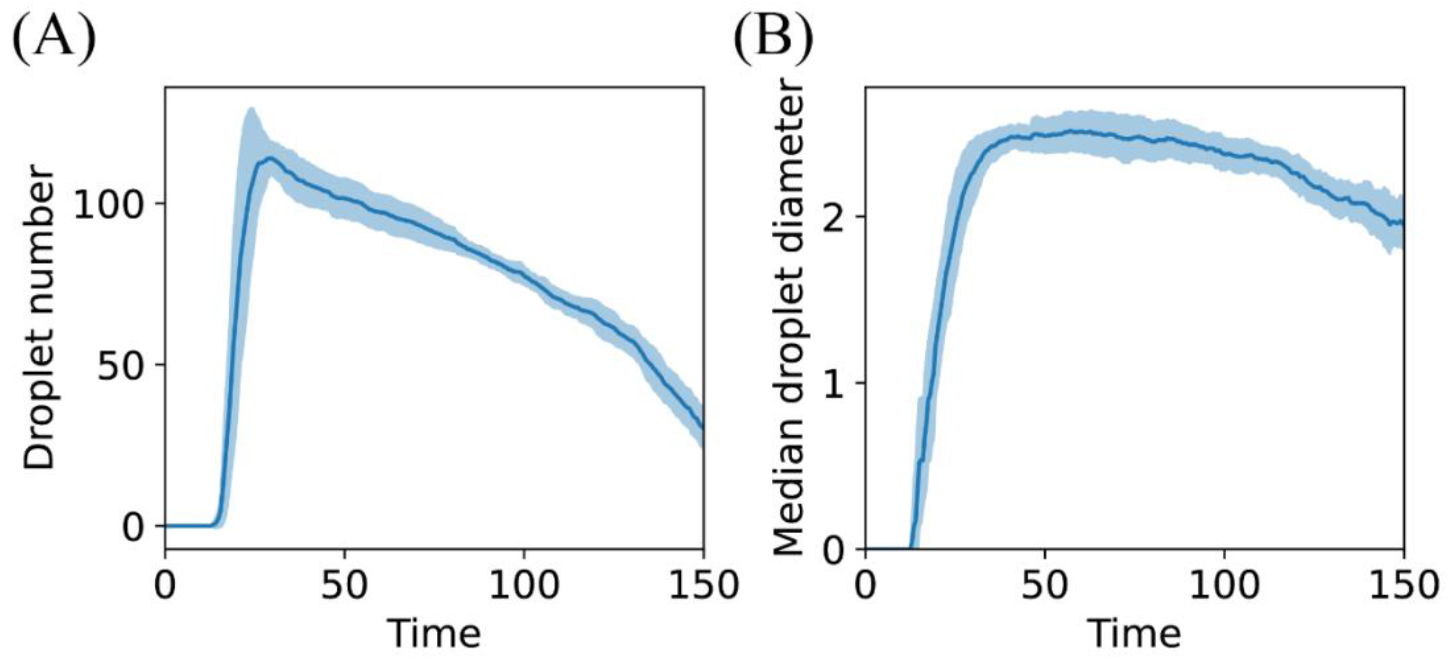
Evolution of 2D droplet population dynamics from the simulation in Fig. 2B. (A) Temporal variation in droplet count. (B) Median droplet diameter as a function of time. Error bars indicate standard deviations (SD) across 10 independent simulation runs. The droplet size is determined by setting the threshold of RAD23B *ϕ*_*R*_ = 0.15, and it is assumed to be circular with its diameter derived from its area.

### 3.2 Transition boundary of proteasomal LLPS

We next investigated the influence of different parameters on the LLPS behavior of the system. The maximum droplet was censused when the LLPS was simulated with tuned parameters such as the initial substrate *ϕ*_*S*0_, the relative changes in *R-S* interaction strengths *Χ*’s and proteasomal degradation rate *k*^(*d*)^ as adopted in Fig. 2. The results are demonstrated in Fig. 4. When the initial substrate *ϕ*_*S*0_ is low, the system fails to undergo LLPS, unless *ϕ*_*S*0_ is increased to exceed a critical concentration threshold, beyond which LLPS emerges (see Fig. 4A and 4B). This observation indicates the existence of a well-defined transition boundary for proteasomal LLPS. This phenomenon aligns with the previously proposed concept of UPS overload [24], where the LLPS transition boundary effectively demarcates the overloaded and non-overloaded states of the UPS. The transition boundary is also modulated by the *R-S* interaction strengths *Χ*’s (Fig. 4A, 4B). Weakening the *R-S* interaction strength shifts the boundary toward higher initial volume faction *ϕ*_*S*0_, indicating that diminished *R-S* interactions enhance the UPS capacity to process excessive substrate loads and significantly inhibit the formation of LLPS. In Fig.4C and 4D, decreasing the proteasomal degradation rate *k*^(*d*)^ initially shifts the transition boundary toward higher initial *S* fraction *ϕ*_*S*0_ (Fig. 4C and 4D). As the degradation rate approaches zero, the transition boundary moves back toward lower *ϕ*_*S*0_. The simulations of the influence of *R-S* interaction strength and proteasomal degradation rate on LLPS also aligns with experimental findings, where a mutation of RAD23B that weaken its interaction with substrate inhibit the formation of LLPS, while proteasome inhibitors can prevent the disappearance of LLPS [19]. Simulation results also indicated that a modest decrease in the proteasomal degradation rate (*k*^(*d*)^) promotes accumulation of *PS*_1_ and *PS*_2_. This accumulation weakens the average *R-S* interaction strengths (*Χ* ‘s), thereby suppressing LLPS formation (data not shown). However, under negligible translocation-degradation rates (*k*^(*d*)^→0), *PS*_1_ and *PS*_2_ accumulation is diminished due to kinetically limited conversion from *PS*_0_ to *PS*_1_. Under these conditions, *PS*_0_ accumulates dominantly, promoting LLPS nucleation (data not shown).

**Fig. 4.**
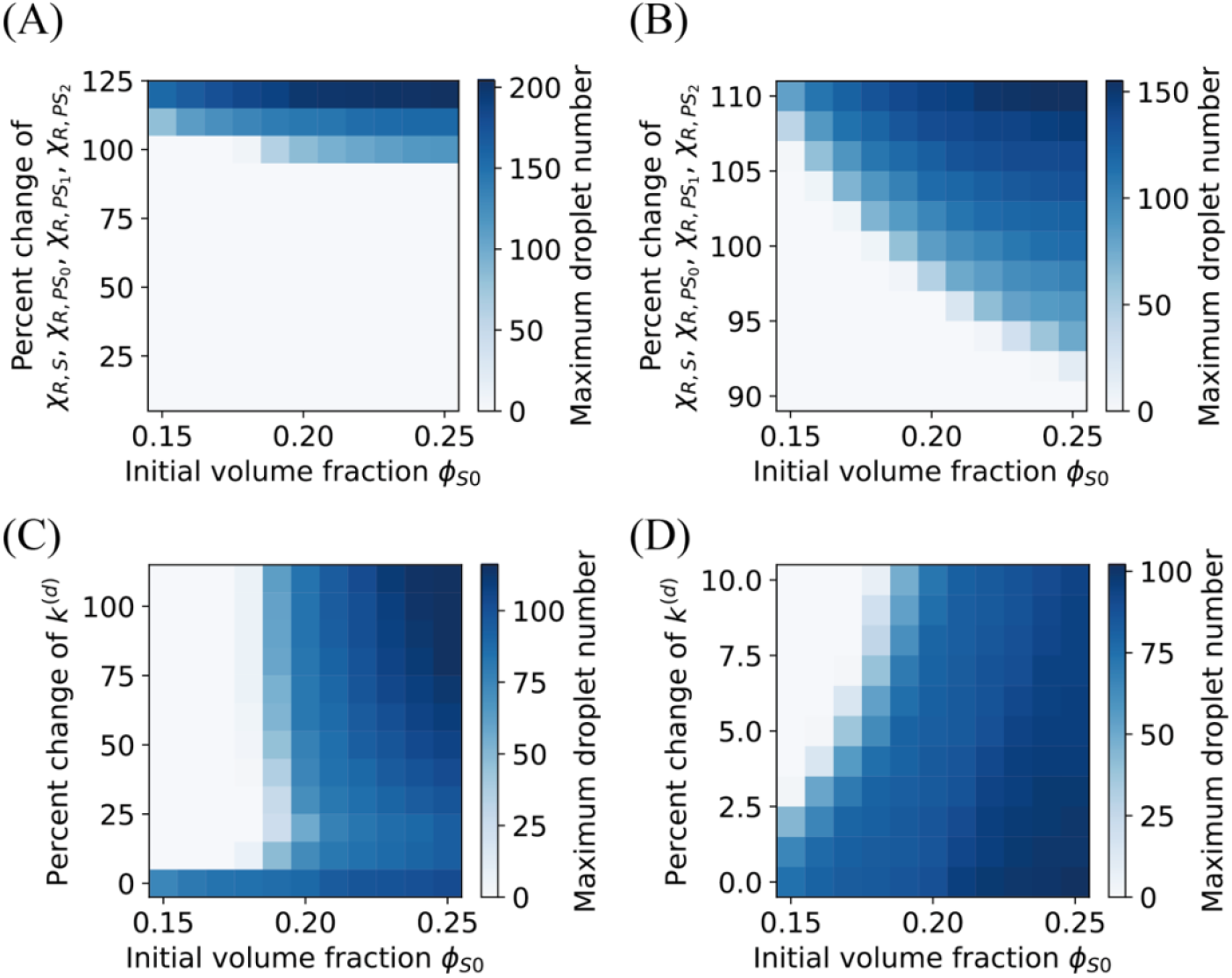
LLPS condensation dependence on system parameters. (A) Maximum droplet count as a function of initial amount of substrate *ϕ*_*S*0_ and relative changes in interaction strengths 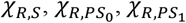 and 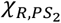 as used in Fig. 2. (C) Maximum droplet count as a function of *ϕ*_*S*0_ and relative change in proteasomal degradation rate *k*^(*d*)^ used in Fig. 2. (B) and (D) are replots of (A) and (C), respectively, both with adjusted vertical axis ranges.

### 3.3 Impact of LLPS on proteasomal degradation

To investigate the impact of LLPS on proteasomal substrate degradation, we monitored the temporal changes in residual substrate volume fraction *S* in situations with and without RAD23B (Fig. 5A). RAD23B depletion abolished LLPS condensate formation, consistent with Fig. 4 showing that weakened R-S interactions disrupt LLPS. In the presence of RAD23B-mediated LLPS (red curve, Fig. 5A), S remained significantly lower than in the LLPS-absent system (green curve, Fig. 5A), indicating enhanced proteasomal degradation via LLPS. Parameter space analysis further revealed that RAD23B-mediated LLPS exerts bidirectional effects on substrate degradation. As shown in Fig. 5B, within the *k*^(*d*)^*-k*^(*b*)^ parameter space, LLPS enhances degradation in regions with high 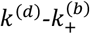 (degradation rate constant) and low 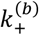 (binding rate constant); LLPS suppresses degradation when 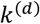 is low and 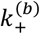 is high. The transition between enhancement/suppression regimes occurs at the boundary.

**Fig. 5.**
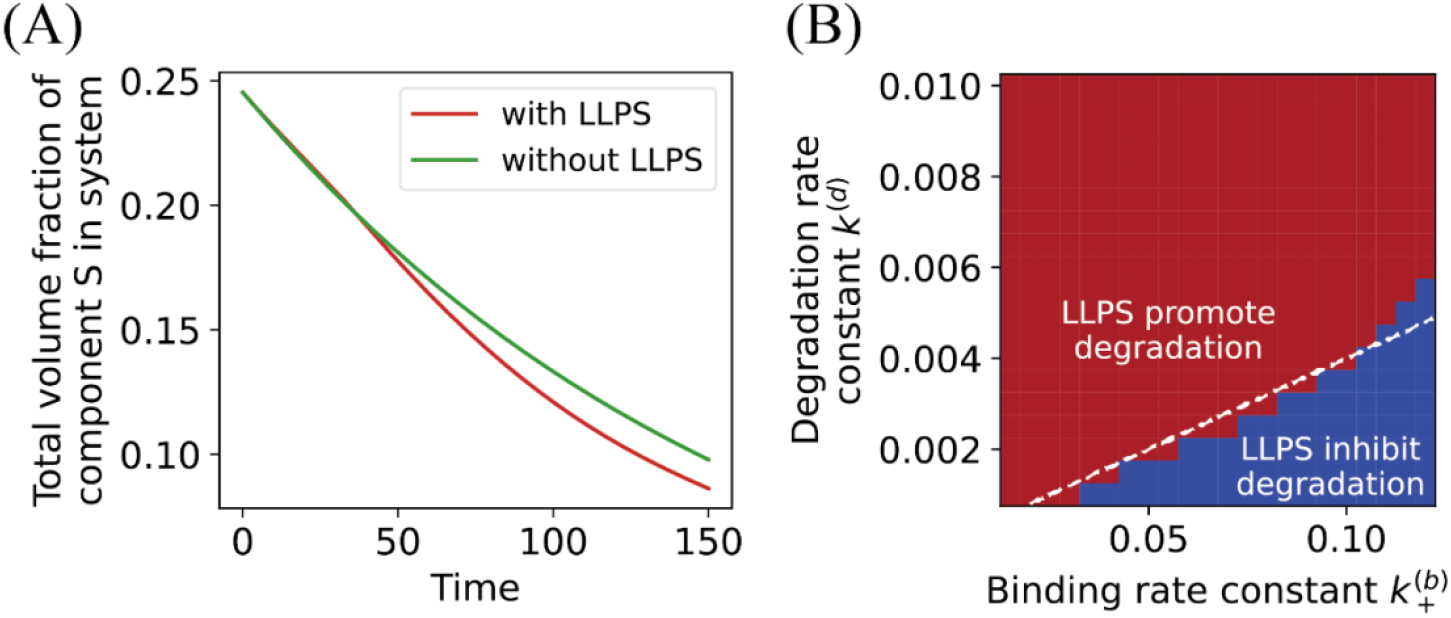
Impact of LLPS on proteasomal degradation. (A) The temporal changes in volume fraction of the residual substrate *ϕ*_*S*_ in the presence (red curve) and absence (green curve) of LLPS. (B) Bidirectional modulation of substrate degradation efficiency by LLPS, dependent on the substrate binding rate constant 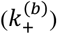 and the proteasomal degradation rate constant (*k*^(*d*)^). The red region indicates conditions where LLPS enhances substrate degradation efficiency, while the blue region denotes conditions where LLPS inhibits degradation. The dashed line represents the theoretical boundary Eq. (11) 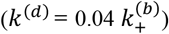 of neutral LLPS effects separating enhancement and inhibition regimes.

This dual effect arises because LLPS differentially modulates substrate binding to the proteasome versus subsequent translocation and degradation. Specifically, LLPS enhances the local concentrations of both proteasomes and substrates within condensates, promoting binding kinetics. Conversely, *R-S* interaction-dependent LLPS formation stabilizes substrates within the condensed phase, thereby impeding proteasome-mediated substrate translocation. The net degradation rate emerges from the competition between the enhanced binding kinetics and impaired translocation. The critical boundary distinguishing the bidirectional effects of LLPS can be derived as follows. Under non-LLPS conditions, the mean substrate degradation time *T* is given by: *T* = *T*_*b*_ + *T*_*d*_, where *T*_b_ and *T*_d_ represent the characteristic times for substrate-proteasome binding and substrate translocation-degradation, respectively. Upon LLPS induction, the binding time is reduced by a scaling factor *α*_*b*_ < 1, while the translocation-degradation time is increased by *α*_*d*_ > 1. Consequently, the modified degradation time becomes, *T*^′^ = *α*_*b*_*T*_*b*_ + *α*_*d*_*T*_*d*_. The condition for net degradation acceleration by LLPS requires *T* > *T*^′^. As the substrate-proteasome dissociation is negligible, the characteristic times are 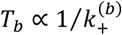, and *T* ∝ 1/*k*^(*d*)^. From *T* > *T*^′^, one has,

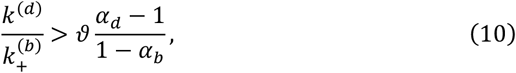

where ϑ is a coefficient. By assuming constant scaling factors *α*_*b*_ and *α*_*d*_ across varying 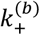 and *k*^(*d*)^, the boundary between LLPS-enhanced and LLPS-suppressed degradation in the 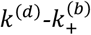 space is a linear relation,

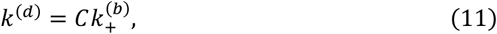

where *C* is a constant. The dashed line in Fig. 5B delineates the boundary between LLPS-enhanced and LLPS-suppressed degradation regimes. This boundary was predicted by the theoretical relation Eq. (11), which shows close agreement with the simulation result.

## 4 Discussion

In recent years, the phenomenon of LLPS in biological systems has garnered increasing attention. LLPS participates in critical physiological processes such as chromatin organization, DNA damage response, DNA transcription, and signal transduction [29]. It not only plays a vital role in normal cellular physiological activities but is also closely associated with the development of diseases such as cancer [30]. Understanding the formation and regulatory mechanisms of LLPS from both experimental and theoretical perspectives is crucial for elucidating its biological significance. The involvement of proteasomes in LLPS introduces an additional layer of complexity to the understanding of proteasome dynamics and the UPS. To quantitatively elucidate proteasome-involved LLPS and uncover its biological functions, we developed an eight-component reaction-diffusion model based on experimental evidence to describe the underlying biological processes. The model recapitulates the experiment observation of droplet formation followed by disappearance, while demonstrating qualitative agreement with experimental findings.

In the modeling, we primarily referenced the RAD23B-mediated LLPS described in [19]. The overloaded and non-overloaded states of the UPS are effectively defined by the phase transition boundary of LLPS. These findings may potentially extend to other proteasome-involved LLPS phenomena. For instance, in p62-induced LLPS, the interactions between p62 and other components not only resemble those of RAD23B, but the emergence and disappearance of droplets are also similar with RAD23B-induced LLPS [21,22]. This suggests that p62-induced LLPS may share identical mechanisms with RAD23B-induced LLPS, differing only in kinetic parameters between the two systems. Whether our model can be generalized to explain other types of LLPS with proteasome requires further experimental validation.

Although there is no clear evidence that LLPS can promote or inhibit the degradation of substrates by the proteasome in RAD23B-mediated proteasomal LLPS, studies have shown that LLPS accelerates the degradation of substrates by the proteasome in p62-mediated proteasomal LLPS [21]. Our computational results indicate that LLPS can not only promote but also suppress substrate degradation by the proteasome, a prediction not yet observed experimentally. Previous studies on the autophagy system have demonstrated its functional interplay with UPS [24,31]. When UPS overload results in substrate aggregation, the autophagy system can eliminate the aggregates through autophagosome encapsulation and subsequent lysosomal degradation. Our findings suggest that when LLPS inhibits UPS function, the autophagy system serves as a necessary complement to UPS for substrate degradation. Our work provides a research foundation for investigating the mutual regulation between the UPS and autophagy systems linking through LLPS and constituting a larger-scale dynamic system.

## Data availability

The source code and data used to produce the results and analyses in this manuscript are available on GitHub at https://github.com/diwu31415/proteasome_LLPS.

## Acknowledgments

This work was supported in part by the National Natural Science Foundation of China (Grant No. 12090051), and the Starry Night Science Fund of Zhejiang University Shanghai Institute for Advanced Study.

